# Information seeking and simulation: Roles of attention in guiding a goal-directed behavior

**DOI:** 10.1101/104091

**Authors:** Rei Akaishi, Eiji Hoshi

## Abstract

We usually actively seek out the information we need. However, it is still debated whether information seeking in decision situation is a purposeful behavior or a random process. We investigated this issue using the decision task involving multiple goal-directed event sequences, in which a contextual cue specifies an associated target and touch to the target delivers the reward. We found that the gaze followed the sequence of contextual cue to the associated target, which was eventually chosen. This fixation sequence from contextual cue to the associated target could be observed even when there were multiple goals and when the focus was shifted from one goal to another. To causally investigate the effects of sequential simulation, we directly manipulated the processing of the contextual cues and found its influence on the final choice of target. Furthermore, past episodes of the sequences influenced both final choices of targets and initial gaze to contextual cues. We interpret the results as suggesting that the internal process of simulating goal-directed event sequence drives information-seeking behavior such as attention/gaze in decision situations.

## Introduction

When we consume information, we are rarely passive recipient. Rather, we actually seek out the information we desired. For example, we click and move to the specific webpage with some expectation of the content and interpret the information with that prior conception. Thus, when we seek new information, somewhat paradoxically, we already have some knowledge of its content. However, it is still debated how and why the prior knowledge can influence the selection of the information to be processed.

Such decision to choose and process specific information has been studied in the research of attention^1,2^. There have been proposals about the factors guiding attention. One line of thought is purely informational: information seeking behavior is to satisfy the pure need for information, which in turn determined by the gap between current level of the knowledge and the information available in the environment^3,4^. Another line of thought is that information seeking behavior is driven by valence or reward-related aspects of the information^1,4–8^. These two types of the influences on information seeking behavior are treated separately^1,4,5^. However, recent studies suggest that there are interactions between the two factors^4,9,10^. There can be some mechanism mediating the interaction between the informational and reward-related factors or there can be an intrinsic link between the two factors.

Attention is also known to guide the direction of our action to achieve a goal. For example, we always pay attention to the direction we are heading when we are driving a car^11,12^. If you intend to turn right, your gaze scans the rightward spaces of for possible hazards and collisions. In more complex situation involving series of steps such as cooking, attention follows the sequences of objects and places where steps of goal-directed behavior are likely to occur^13,14^. In a goal-directed behavior, actions are executed in sequence and there is sequential dependency between these successively occurring events^11,14,15^. It is possible that the internal mechanisms of the brain might exploit such sequential dependency in natural behavior. More precisely, the events in sequence can be represented as a chunk of actions or events^16–20^. It is also reported that the neurons in hippocampus shows the pattern of neural activity as if the animal is simulating the sequences of the future behavior toward the reward/goal position ^21–23^. There have also been attempts of incorporating such sequential nature of event structure in model-based approaches of reinforcement learning^24–27^. Thus, it is possible that attention is an intrinsic part of the mechanism to execute a goal-directed behavior.

Our proposal is that simulation of these sequential events might drive the attention and gaze through the sequence of objects and places leading to the goal/reward. In turn, the attention to each step of the sequence can reduce uncertainty inherent in the variations of the events in unique episodes. In this framework, it is natural to observe that the attention/gaze is attracted to the events/places associated with the reward and also works to reduce uncertainty about the individual events for smooth execution of goal-directed behavior. This simulation-based account of attention is consistent with the recent results reporting the interactions of information seeking and reward/valence aspect of the information^4^. One study reported that information seeking behavior of humans in card selection task was biased toward more valuable option^9^. Even in monkeys, information seeking behavior with gaze response showed the preference for the advance information of the options with higher rewarding outcome^10^. In other words, the subjects sought after or pay attention to the information about the specific option or course of action leading to more desirable outcome.

However, it is still not known how simulation-based mechanism of attention works in the decision situations where multiple goals can compete for attention or more weights in information processing. The issue is that choices in a realistic situation entail selection of one of the multiple sequences of stimulus/action events leading to different goals. In a simple choice task, it has been suggested that randomly directed attention/gaze the options sample the information from these options and these sampled information are accumulated to make a decision^28–30^. However, it was recently reported that the direction of gaze of human subjects to options is not so random and might be related to the simulation in sequential multi-step decision tasks^31,32^. In other words, the gaze directed to the options in the decision situations might not be random process sampling decision-related information. Rather, as in other situations of information seeking and goal-directed behavior, a decision maker would actively use the prior knowledge about options by simulating events associated with each option.

We investigated this issue using the task design involving an event sequence in which a contextual cue specifies an associated target and touch to the target delivers the reward. We found that the eye movements in this task followed the sequence of the associated events leading to the goal/reward. Importantly, the sequence following eye movements could be observed even in the choice situation with the multiple goal-directed sequences of contextual cues and corresponding associated targets. Furthermore, these patterns of eye movements following associated sequence could be observed even when the final target choice is changed from the initially followed sequence. To causally test the relationship between sequence simulation and goal selection, we manipulated the initial processing of the contextual cue by priming and found that the enhancing the processing of the contextual cue influenced the final choice of the target. We also examined how the prior knowledge was used for sequence simulation and selection by analyzing the effects of past episodes of the event sequences. We found that the past episodes influenced both the initial gaze to contextual cues and the final choice.

## Results

### Task and behavior

To test the sequential operation of the goal-directed behavior, the task was structured with the sequence of contextual cue, target, target touch and reward in this order. A specific contextual cue determined which of the two targets would deliver reward when touched (Fig. 1A).

**Figure 1.**
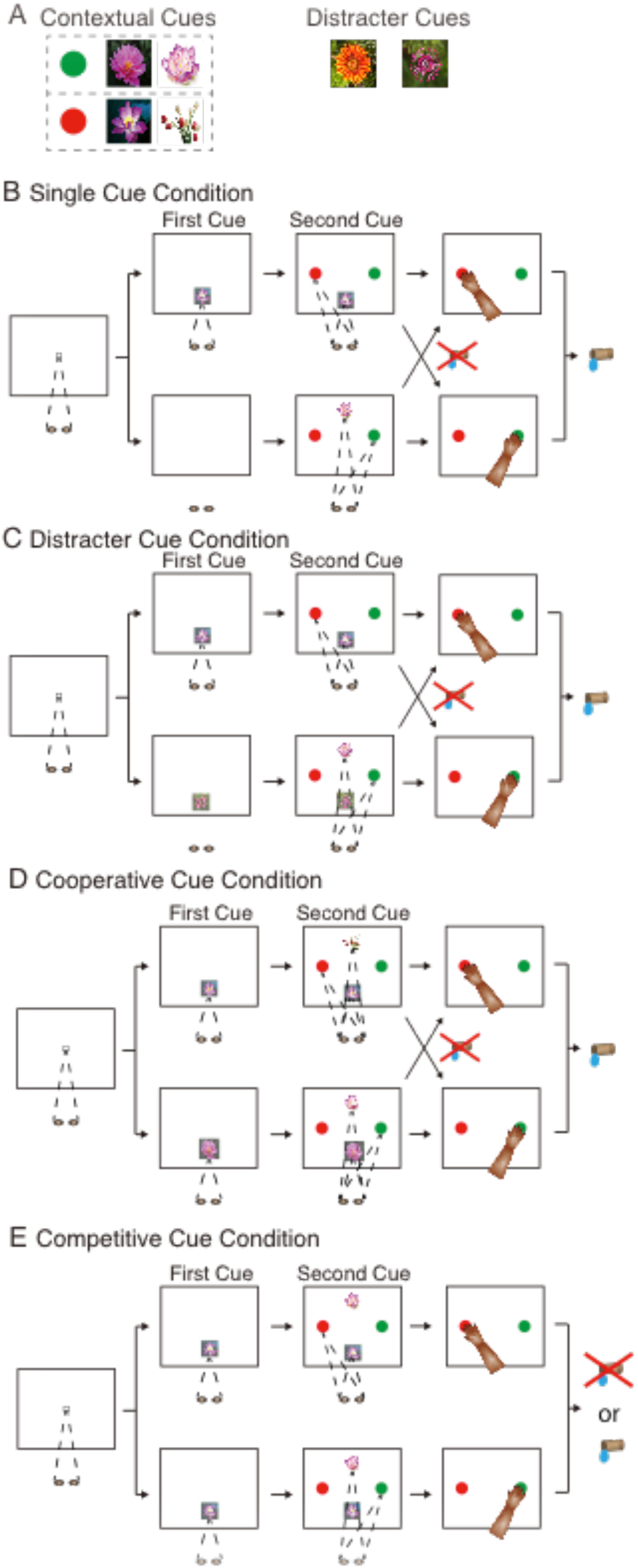
Task design and gaze behavior. (A) Four pictures of flowers were used as contextual cues and each of them was associated with a specific target (green or red). The touch of the associated target delivered the reward. The two other cues were not associated with any target and these cues did not predict which target led to reward delivery when touched. (B) Single cue condition. A press of hold button triggered appearance of the fixation on the screen. Sequence of the upper branch: After the gaze at the fixation for 500 ms, the first contextual cue appeared. The monkey could freely move their eyes thereafter. After the period of 300 ms, the two targets appeared. The monkey could then initiate reaching by releasing the hold button, which extinguished the contextual cues. Finally, the monkey touched a target. If an appropriate target was touched, a juice reward was delivered. Sequence of the lower branch: The contextual cue could appear at the same time with the target. See more details in the text. (C) Distractor cue condition. The task sequence is same as Single cue condition except that the distractor cue appeared at the timing the contextual cue is not presented. Distractor cues did not instruct the correct target to touch for obtaining the reward so that the subjects had to rely on the contextual cue to determine the rewarding target. See more details in the text. (D) Cooperative cue condition. The two contextual cues were presented but both of them instructed the subjects to touch the same target to be rewarded. So, whether the subjects used the first cue or second cue, the same target could be chosen as rewarding target. Either the red target (upper branch) or the green target (lower branch) could be chosen based on the association with the contextual cues. See more details in the text. (E) Competitive cue condition. The two contextual cues were associated with the different targets so that the use of the different cue could result in the choice of the different target. For example, the use of the first cue resulted in the choice of the red target (upper branch). In contrast, the use of the second cue led to the choice of the green target (lower branch). The order, position, the association with the targets were pseudo-randomized in the actual experiments. See the text for more details.

Depending on the specific combinations of the cues, the conditions were divided into four types: Single cue condition, Distractor cue condition, Cooperative cue condition, and Competitive cue condition.

#### Single cue condition

In a given trial, the monkey subject first pressed a hold button and the fixation cue appeared in the center of the screen (Fig. 1B). The subject then fixated on it for 500 ms and the first cue appeared in a half of trials (upper branch of the sequence). After 300 ms of waiting period, the two targets (red and green) were presented at 90-degrees-rotated positions. These arrangements of the contextual cues and targets were exchanged across blocks of 10-20 trials. The monkey released the hold button triggering extinction of contextual cue(s) and touched the target. Touching the rewarding target, which was associated with specific contextual cues (Fig. 1A), triggered delivery of a juice reward. A single contextual cue associated with a specific target determined the rewarding target when it was presented alone in Single cue condition. The contextual cue can be presented either at the pre-target timing (first cue; upper branch of task sequence in Fig. 1B) or the same timing as the targets (2nd cue; lower branch of task sequence in Fig. 1B). In the latter case, subjects waited for 800 ms for the cue and targets to appear.

#### Distractor cue condition

Distractor cue condition was same as Single cue condition except that a distractor cue, which was without any association with targets, was presented with a contextual cue (Fig. 1A and C). As in Single cue condition, two targets were presented and the monkey subjects had to touch one of them to obtain the reward according to the association of the presented contextual cue with one of targets. Only one contextual cue was presented in a trial and it was either at the first cue timing (the upper branch of the sequence in Fig. 1C) or at the second cue timing (the lower branch of the sequence). The Distractor cue was also presented either at the first cue timing (the lower branch of the sequence) or at the second cue timing (the upper branch of the sequence), which was the different timing from the contextual cue.

#### Cooperative cue condition

In Cooperative cue condition, the two contextual cues were associated with the same single target (Fig. 1D). So the subjects still need to select one of the two targets but the selection was easy because both contextual cues instructed the touch of the same target. If the contextual cue presented at the first cue timing was associated with the red target, the contextual cue presented at the second cue timing was also associated with the red target (the upper branch of the sequence in Fig. 1D). The two contextual cues can be associated with the green target (the lower branch of the sequence in Fig. 1D).

#### Competitive cue condition

Competitive cue condition involved the two contextual cues, which were associated with the different targets. Thus, both targets could deliver the reward when touched. But the rewards were delivered only 50% of time regardless of choice of touched target. However, the monkeys did not know whether they were in the Competitive condition before the presentation of the second cue. The monkeys usually completed the trials of Competitive cue condition as in other conditions and aborting of a trial was very rare. Because other conditions, which were a majority of the trials, had a single contextual cue and a deterministic rewarding target, their strategy was to rely on one of the contextual cue to choose one of two targets. In one case, the monkey subject relied on the contextual cue presented at the first cue timing and chose one of the targets (a red target in this case) (the upper branch of the sequence in Fig. 1E). The case of using the contextual cue presented at the second cue is described in the lower branch of the sequence (Fig. 1E). This point will be revisited when we analyze this condition in detail.

In total, 40% of all trials were for Single cue condition and 20% of all trials for each of other conditions. Thus, in the most of trials (40% + 20% + 20% = 80%), the contextual cue instructed the monkey to touch the single rewarding target. Probabilities of rewarding choice were high (mean of 90% across conditions) except the competitive condition (50%; Fig. 2A). All the contextual cues and the distractor cues can be presented either at the pre-target timing (1st cue) or the same timing as the targets (2nd cue). The four conditions were pseudo-randomly interleaved across trials and the monkeys could not know which condition they were in before the presentation (or absence) of the second cue.

**Figure 2.**
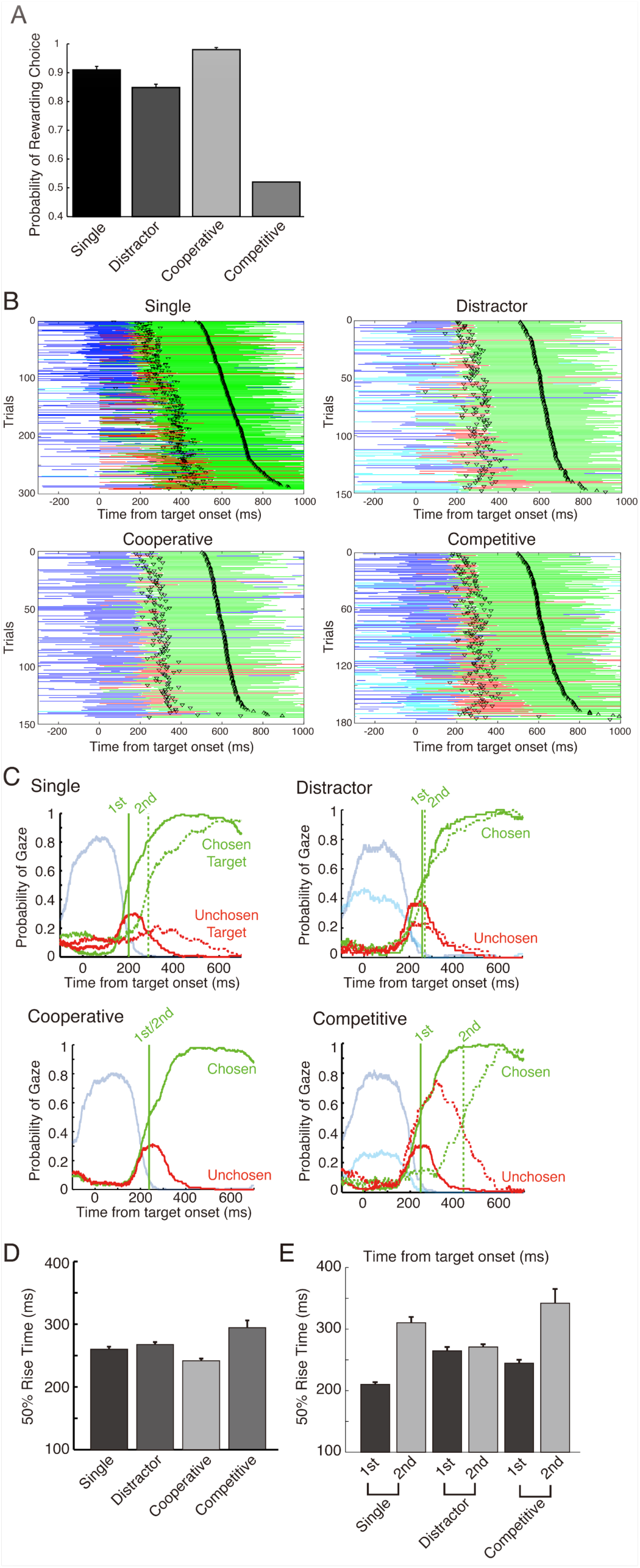
Behavior across task conditions. (A) Probabilities of rewarded choices are plotted for all conditions. For Single, Distractor, and Cooperative cue conditions, the contextual cues presented in a given trial determined the rewarding choice of a target from two alternatives. For Competitive condition, the reward probability is 50% regardless of the choice of the target. The actual probability of the rewarded choice in this condition was at chance level. Data are presented as mean ± s.e. across 17 sessions. (B) Example raster plots of the gaze patterns across trials for four conditions. The time course of the gaze fixations are plotted with distinct colors for different visual objects: green for the touched (chosen) target, red for the not touched (unchosen) target, blue for the contextual cue associated with the chosen target, cyan for the contextual cue not associated with the chosen target. The trials were sorted according to the target touch time (top-pointing triangle, △) from the target/second cue onset (0 ms). The movement initiation times (release of hold button) were also plotted with lower pointing triangle (▽). (C) The gaze likelihood measures for four conditions. The metric at each 1ms-bin was calculated as probabilities of gazes falling on the specific visual objects that were present. The color coding was same as the gaze rater plots in (B). Trials were distinguished based on whether the final target touch was consistent with the cue presented at the first timing (before target onset; solid green and red lines) or the second timing (simultaneous with target onset; dotted green and red lines). Note that probabilities of target fixations sum up to 1 because the two targets (chosen and unchosen) were always presented together. On the other hand, the contextual cues were separately presented at separate timing in different trials and the gaze likelihoods were calculated separately. (D) 50% rise time across four conditions. It is calculated as the time the gaze likelihood reach the half value of the peak probabilities for the first time. Note that Competitive cue condition was taking more time compared to other conditions. (E) 50% rise times were calculated separately for the cases of the choices consistent with the contextual cue presented at the first timing and the second timing. In Single cue condition and Competitive cue condition, these two timings were different.

### Gaze patterns in four conditions

We first examined the qualitative pattern of eye movements in each condition. In a trial, the gaze of the monkey subject was usually first directed to contextual cues (Fig. 2B). This was especially true when the contextual cue was presented at the first cue timing, which was before the onsets of the targets (fixations on the contextual cue associated with chosen target; shown in blue traces before time 0). After the contextual cues were fixated, the gaze was frequently oriented toward the associated action targets (see the relative abundance of the cases of transitions of blue-to-green traces). Notably, the gaze was directed to the target to be touched even almost always before the initiation of the reaching movement (down-pointing triangles, ▽, for the release timing of hold button and top-pointing triangles, △, for timing of target touch in Fig. 2B). Such precedence of the gaze before the hand movement was reported in previous studies^13,33^. However, to our knowledge, the gaze to the cues associated with the motor targets has not been systematically analyzed.

To quantitatively examine the gaze patterns to the contextual cues and targets, we used the gaze likelihood measure^34^. The gaze was first directed to the contextual cue that was associated with the chosen target during the time period from the onset of the contextual cues to the timing shortly after the target onset (Fig. 2C). The gaze was then increasingly directed toward the target to be touched (Chosen Target) where as the gaze to the untouched target (Unchosen target) decreased. The ramping patterns of the gaze likelihood were different across conditions. To quantify the ramping patterns for each condition, we calculated the timing when the gaze likelihood reached the half value of the peak of the gaze likelihood for the first time (50% rise time). Across conditions, 50% rise times are significantly different (p=0.000013). Compared to Single cue condition (260 ms), the 50% rise time of the Distractor cue condition (268 ms) was slightly later (p=0.11) (Fig. 2D; the results were consistent across monkeys, Supplementary Figure 1). The 50% rise time of the Cooperative cue condition (242 ms) was significantly shorter than that of Single cue condition (p=0.000054). 50% rise time of the Competitive cue condition (293 ms) was significantly delayed compared to that of Single cue condition (p=0.0052) probably because of conflicting information from the two cues.

Across conditions, the gaze tended to be directed to the target earlier if it was associated with the contextual cue presented at the first timing (solid green vertical line) compared to the gaze to the target associated with the contextual cues presented at the second timing (dotted vertical green line) (in Cooperative condition, the solid and dotted lines were merged because the contextual cues presented at both timings instructed the touch of the same target) (Fig. 2C). This observation was consistent with the proposal that the gaze is following the sequence of associated events in a goal directed behavior. More precisely, if the initial gaze to the contextual cues signifies the initiation of simulating an association sequence, the gaze to associated targets would follow the gaze to the cue in the same association sequence. This sequential nature of gaze pattern was demonstrated in Single cue condition: the gaze to targets was generally delayed when the single contextual cue was presented at later timing (2nd cue) compared to when it was presented at earlier timing (1st cue) (Fig. 2C). The 50% rise time, which is a measure of this target gaze latency, was longer for the 2nd cue case than the 50 % rise time for the case of 1st cue (t-test: p = 0.00000018; Fig. 2E; the results were consistent across monkeys, Supplementary Figure 1).

The gaze pattern in the Single cue condition was consistent with the idea of simulation of associated sequence. However, it was not clear whether the same process operates in the situation where the multiple possible association sequence exists. In our task design, some trials contain multiple contextual cues each of which the subjects can use to simulate the associative structure relevant for a choice of a specific target. These contextual cues are presented sequentially so that timing as well as the position of the gaze to either target can tell the experimenter which specific association sequence was followed. We focused on Competitive cue condition because the contextual cues were uniquely associated with different targets so that the touch choice of the target could reliably identify the associated contextual cue. The ramping patterns and 50% rise times of cue cases were reliably later when the chosen target was associated with the 2nd cue than the cases when the chosen target was associated with the 1st cue (t-test: p = 0.0011; Fig. 2E; the results were consistent across monkeys, Supplementary Figure 1). We could not conduct the same analysis to trials of Cooperative cue condition because the contextual cues were associated with the same target. We could not observe the clear effect in Distractor cue condition (p > 0.05) probably because the distractor cues could be associated with both targets and which cue was used in the targets selection was not unambiguously identified in the analysis.

### Influences of the gaze to contextual cues on target selection

The gaze patterns across conditions were generally consistent with the idea that attention follows the sequence of associated events in a goal directed behavior. The next question is whether this pattern of attention/gaze can be observed in the situations where multiple goal-directed sequences exist. To test these ideas, we analyzed the gaze to contextual cues in Competitive cue condition in which the two contextual cues were associated with different targets. The monkey subjects could select a target relying on either one of the contextual cues as in the other conditions. Based on the target choice indicated by the touch, we designated the contextual cues as either a choice-consistent (CC) cue, which was associated with the chosen target, or a choice-inconsistent (CI) cue, which was associated with the unchosen target (Fig. 3A).

**Figure 3.**
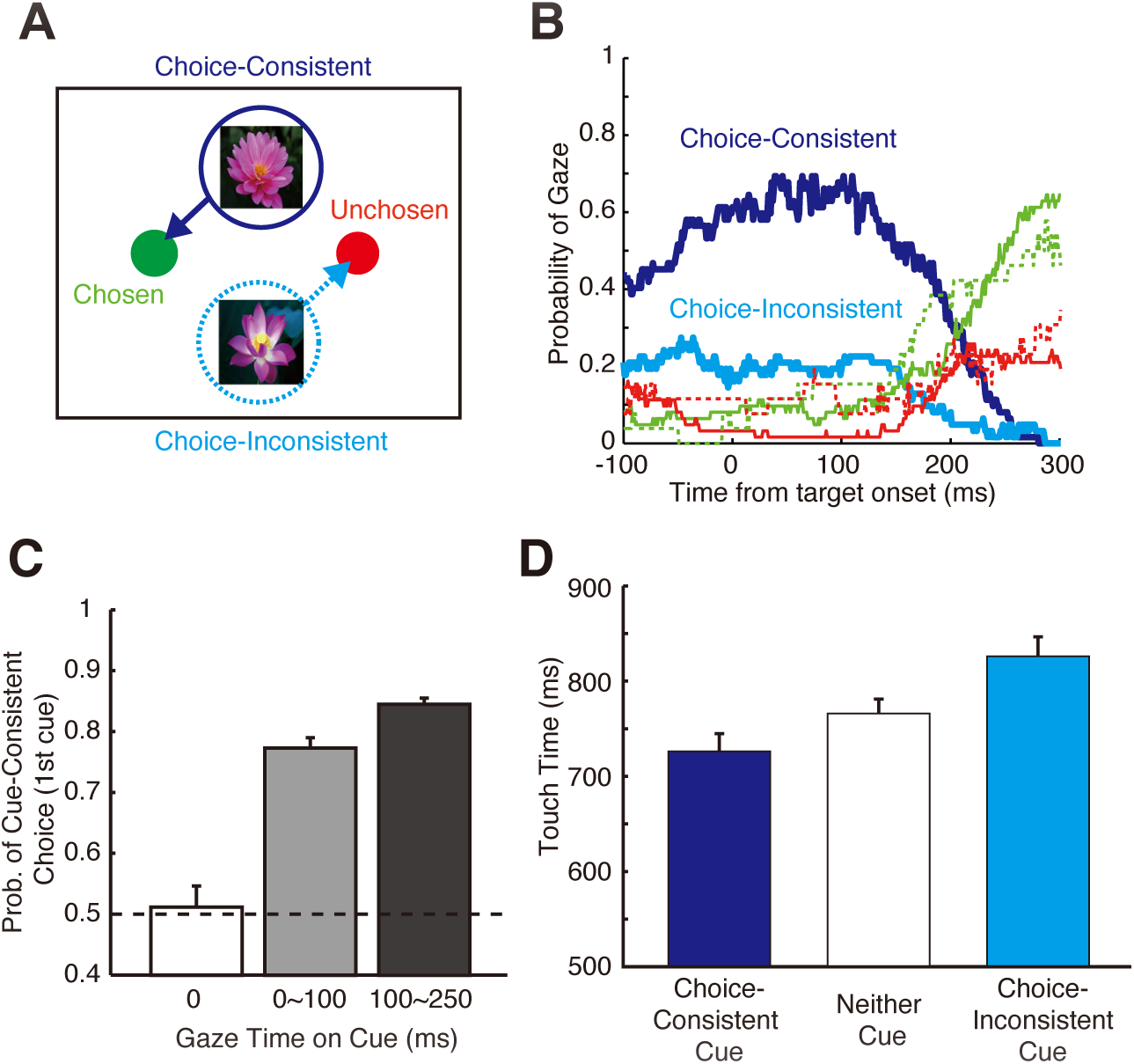
Behavioral effects of gaze to contextual cues. (A) In Competitive condition, choice-consistent (CC) cue (blue) was associated with the chosen target (green) whereas choice-inconsistent (CI) cue (cyan) was associated with the unchosen target (red). (B) The gaze likelihood shows that CC cue (blue line) was fixated with higher probability than CI cue (cyan line) in the period before the gaze to targets (pale green and red lines). (C) If the contextual cue (first cue) was fixated during the 250 ms after target onset, the subsequent choice was more likely to be the associated target. The probability of the cue-consistent cue increased from 75% (Different from 50%: t(16) = 13.35, p = 4.3^−10^) to 84% (Different from 50%: t(16) = 36.8, p = 6.8^-17^) as the time of fixation increased (0~100 ms to 100~250 ms) (paired t-test: t(16) = 4.84, p = 0.00018). On the other hand, it was at the chance level o (51.1%; t(16) = 0.33, p = 0.75) if the contextual cue was not fixated. (D) We examined touch time, the duration from the target onset to target touch. Compared to the case when neither cue was fixated (766 ms), the visual fixations (> 100 ms) on CC cue shortened the touch time (726 ms). In contrast, the fixation on CI cue lengthened the touch time (826 ms) (paired t-test: t(16) = -3.51, p = 0.0029; t(16) = 4.29, p = 0.00056). Data are presented as mean ± s.e. across 17 sessions.

If attention in decision situation is not a random process but reflects on the underlying simulation process, the gaze would differentiate the CC cues, which was in the same simulated sequence with the chosen target, from the CI cues, which was not in the same sequence. We first analyzed the pattern in the gaze likelihood measure ^34–36^, to examine the differentiation of the cues contingent (CC cue) or not contingent (CI cue) with a final choice of a target. Monkeys had more visual fixations on the CC cue than the CI cue (Fig. 3B; also see Fig. 2B and C for trial-by-trial patterns and data from other conditions). This is consistent with our hypothesis that the target selection starts with the engagement with a contextual cue in a specific event sequence. Note any bias of selecting the first cue does not contribute to this result because the first and second cues were almost equally likely associated with the final choice: the first cue was associated with the chosen target for 57% of times.

It is important to ask how these results are related to the previously observed effects of fixations on options^28–30^. To precisely examine how the initial fixations on the contextual cue were related to a subsequent choice of target, we analyzed the probability of the choice consistent with the first cue depending on how much the gaze was fixated on it. The probability of a choosing the target consistent with the first cue was just 51.1 % if the monkey subject did not look at it (Fig. 3D). In contrast, if the monkey took a cursory look at the cue with a short duration (fixation time < 100 ms) ^37^, it produced cue-consistent target choice at 75.0 %. With longer fixations (fixation time > 100 ms), the probability of the cue-consistent choice is higher (84.0 %). Thus, the influence on the target choice was dependent on the time length of the gaze to the associated cues in a specific sequence (the results were consistent across monkeys, Supplementary Figure 2). We also found that the visual fixations on the contextual cues modulated time between target presentation and target touch (Fig. 3E). Compared to the case when the subject did not look at any first cue (766 ms), the fixations on the CC cues shortened the touch times (726 ms; paired t-test: p = 0.00056; the results were consistent across monkeys, Supplementary Figure 2) whereas the fixations on the CI cues (at least 100 ms) resulted in longer touch times (826 ms) (paired t-test: p = 0.0029).

These influences of gaze on the contextual cues can be interpreted as a pure manifestation of information sampling as reported in the previous studies^28–30^. However, the gaze behavior and information seeking can be interpreted as a reflection of simulation process ^9,10,31,38^. Specifically, these results can indicate that the simulation of association sequence made it easier to choose the associated target in that sequence and difficult to choose the target not in the simulated sequence. This is an equally plausible account given that we have also observed the results consistent with the explanation of simulation-based mechanism of attention (Fig. 2). We will contrast these alternative accounts in the subsequent analysis of sequential gaze patterns from the contextual cues to targets.

### Sequential gaze patterns

If the gaze sequence from contextual cues to the associated targets reflected the process of simulating the association sequence, it must also show the influences of these simulation processes regardless of ultimate choice. The simulation process reflected in the target gaze might also exhibit the momentary focus on the specific cue-targets association sequence even in the cases when the simulation process switched from one association sequence to another. Such switching mechanism is expected in natural behavior, which requires tracking of multiple operations such as driving and cooking^11^. In contrast, random process of information sampling would not predict the strong sequential pattern of gaze from the contextual cue to the target. Especially, in the case when the final choice was inconsistent with the initially fixated contextual cue, the gaze would favor the target to which more evidence for preference is supposedly accumulated^28,29,34^.

To analyze the sequential gaze patterns, we examined the gaze likelihood to targets separately for the trials when the fixated first cue was associated with the final choice of target (CC cue) or not (CI cue) in Competitive cue condition. In a representative session (Fig. 4A), after the first fixation was directed on the CC cue, the gaze was immediately directed to the chosen target. This was reflected in the corresponding trajectory of gaze probability (green line). Although this is consistent with the our hypothesis of sequential simulation, more critical test of the hypothesis was whether the gaze to the associated target followed immediately after the initial fixation on the CI cue. As predicted by the hypothesis, the gaze was directed to the unchosen target soon after the fixation on the CI cue (orange line) as if the subject was momentarily considering the possible association sequence from CI cue to the associated but unchosen target. The targets that were not associated with the initially fixated cues (cyan and red lines) did not attract the gaze in the period immediately after the initial fixation on the cues.

**Figure 4.**
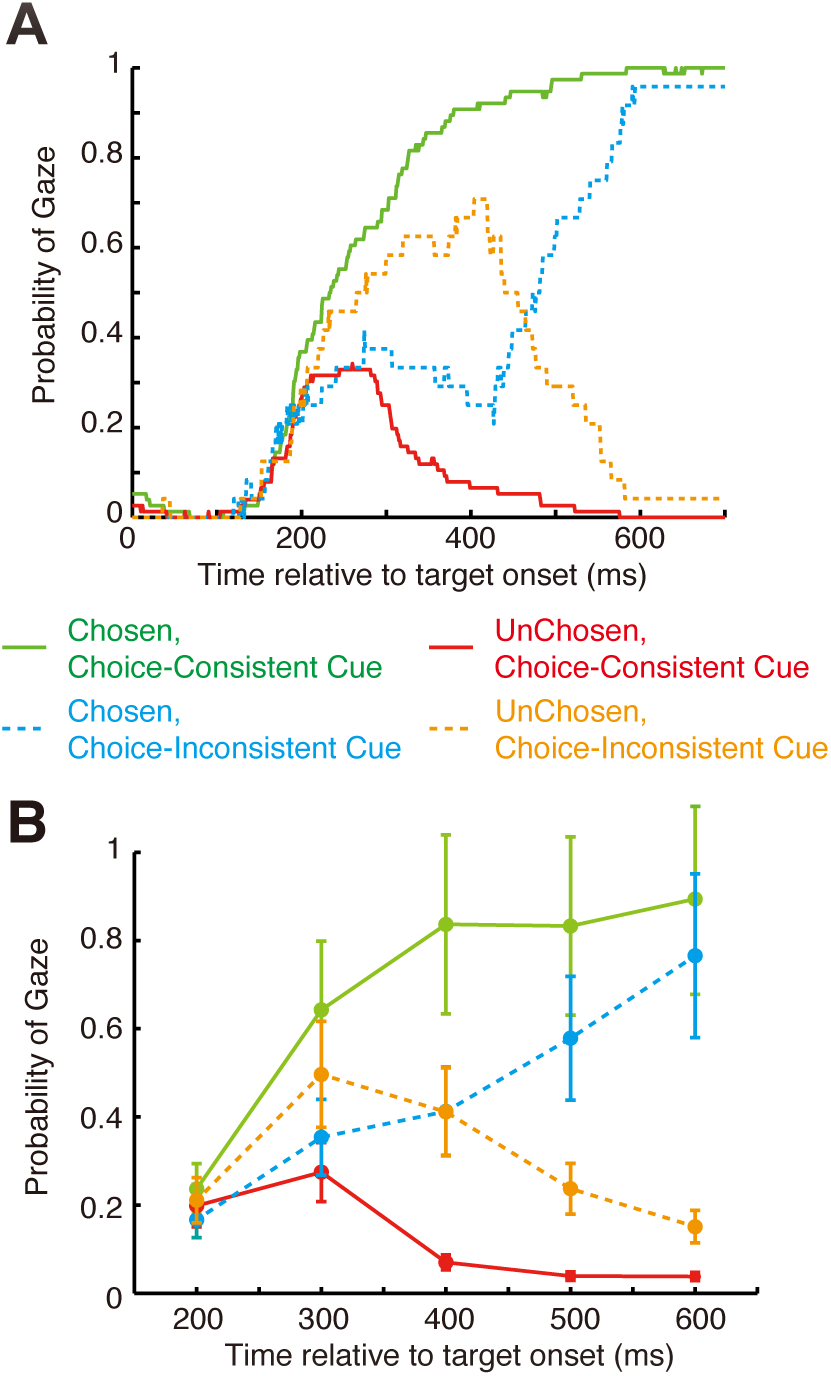
Dependency of target gaze likelihood pattern on initial cue fixation. (A) In a sample session, the gaze to CI cue was accompanied by the greater increase of the gaze likelihood for the unchosen target (dotted orange), which was associated with CI cue, compared to the case with the preceding gaze to CC cue (solid red). The gaze likelihood to the chosen target smoothly rose in the case when the associated cue (CC cue) had been looked (solid green) whereas the gaze to CI cue resulted in the dip of the gaze probability (dotted cyan). (B) The data across sessions showed that from 300 ms after the target onset, the gaze to CC cue produced higher likelihood of the gaze to the chosen target (solid green) compared to the case of the preceding gaze to the CI cue (t-test: 300 ms, p = 4.4^−5^;400 ms, p = 1.2^−8^; 500 ms, p = 8.5^−5^; 600 ms, p = 0.024). On the other hand, the gaze to CI cue resulted in the higher gaze likelihood for the gaze to unchosen targets (dotted orange) compared to the case with the preceding gaze to CC cue (solid red) (t-test: 300 ms, p = 0.0013; 400 ms, p = 1.2^−6^; 500 ms, p = 1.8^−6^; 600 ms, p = 0.018). Data are presented as mean ± s.e. across 17 sessions.

The gaze likelihood data also showed the similar pattern (Fig. 4B; the results were consistent across monkeys, Supplementary Figure 3). From 300 ms after the target onset, the likelihood of the gaze to the chosen targets after the initial fixation on the CC cue (cyan line) was higher than the gaze likelihoods of the chosen target after the initial fixation on the CI cue case (green line) (t-test: 300 ms, p = 4.4^−5^; 400 ms, p = 1.2^−8^; 500 ms, p = 8.5^−5^; 600 ms, p = 0.024). Thus, the gaze to choice targets depended on the previously fixated contextual cues. Similarly, the gaze likelihoods for the unchosen target (orange lines) after the initial fixation on the CI cue was higher than the gaze likelihoods of the unchosen target after the fixation on the CC cue (red line) (t-test: 300 ms, p = 0.0013; 400 ms, p = 1.2^−6^; 500 ms, p = 1.8^−6^; 600 ms, p = 0.018). Thus, the gaze to the target is dependent on the association with the initially fixated contextual cues independent of whether the target is eventually chosen or not.

### Priming sequential simulation

Traditionally, attention has been thought to operate on the individual objects or positions in space. Experimental manipulations of attention have been generally focused on single entities (with some notable exceptions^39^). However, we regard attention as operating with sequential representations. This is the idea we used in examining the data for aforementioned results of attention in a goal-directed behavior. But it is critical to test this idea in a more causal manner. If attention were operating on isolated events independent of other events without any influence of underlying association structure, the manipulating information processing of contextual cue alone would not produce any bias of target choice. In contrast, the sequential simulation hypothesis predicts that the manipulation of the information processing of the contextual cues alone should be able to bias the choice to the associated target because it can initiate the simulation of the associative sequence or enhance the representation of the associative sequence in the decision process.

To selectively enhance the processing of the contextual cues but not the action target, we conducted priming experiments, in which the contextual cues were primed by the flashing stimulus presented 100 ms before the appearance of the contextual cues (Fig. 5A; see **Methods**). This was meant to attract the initial focus of attention to the position where the first contextual cue was to appear. The cue priming produced effects similar to those of the gaze to the contextual cues (Fig. 5B). The priming of the first cue resulted in more choices consistent with the primed cue (planned t-test: p = 0.005; the results were consistent across monkeys, Supplementary Figure 4). The touch times of the choice consistent with the primed first cues were shorter than those of the choices inconsistent with the primed first cue (Fig. 5C; planned t-test: p = 0.008; the results were consistent across monkeys, Supplementary Figure 4). The priming of the second cue did not produce the similar effects probably because attention or processing resources were already directed to the already presented first cue and/or simultaneously presented targets.

**Figure 5.**
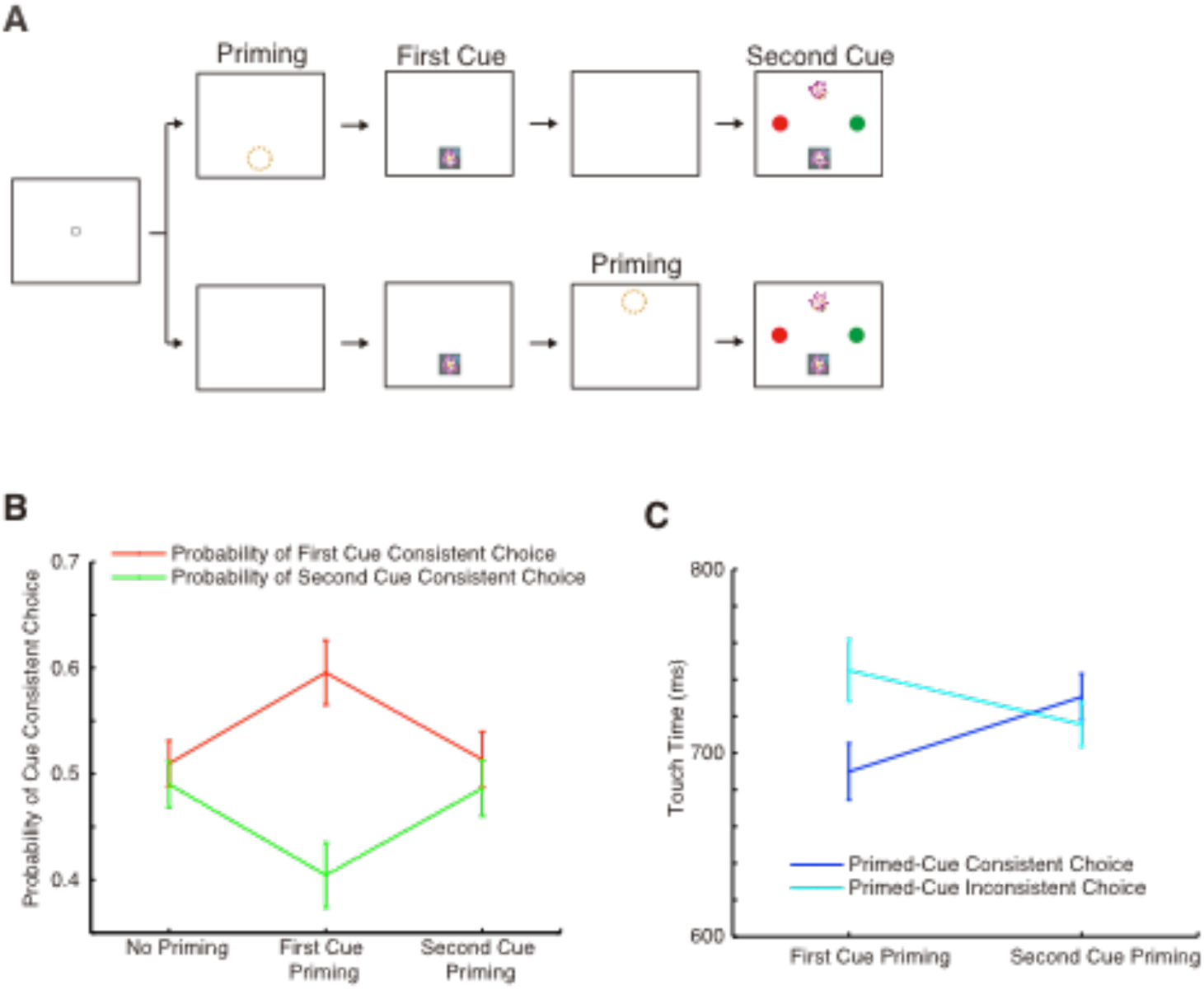
Priming of contextual cues. (A) In the priming experiments, the processing of contextual cues were enhanced by presenting the flashing stimuli at the position of the cues 100 ms prior to the actual cue presentation The upper branch of task sequence depicts the priming for the contextual cue presented at the first timing whereas the lower branch represents the task sequence for the trial of priming the contextual presented at the second timing. The priming was timed according to the presentation timing of the first cue and second cue. (B) We examined the effects of the priming on the probability of the cue-consistent target choice. In contrast to the trials without the priming of any cue (planned t-test: p = 0.66), the priming of the first cue increased the probability of the first-cue consistent choice in competitive trials in the disadvantage of the second cue (planned t-test: p = 0.005). The priming effects were absent if the second cue was primed (planned t-test: p = 0.61). This was probably due to the fact that the spatial attention had been captured by the action targets, which were attracting the gaze at the same time. Then, there were no additional room of processing resources that can be modulated by the priming. (C) The analysis of the effects of the priming on the touch time was also conducted. Consistent with the results of the priming on the cue-consistent choice, the modulation of the touch time appeared in the priming of the first cue (planned t-test: p = 0.008) but not in the priming of the second cue (planned t-test: p = 0.23). Data are plotted as mean ± s.e. across 21 sessions.

These results were generally consistent with the sequential simulation hypothesis and supported the view that the priming of contextual cues enhanced the simulation of the associative sequence so that the choice of the target is biased in favor for the specific target in the sequence.

### Influences of past episodes on gaze and choice

We have suggested that pre-existing knowledge influences information seeking behavior such as attention. Therefore, it is possible that the selection of the simulated sequence would be affected by the past experiences of the same event sequences. This would be also the sign that the monkeys did not acquire the completely habitual mode of operation but could flexibly use the stored information of the event sequences in planning a goal-directed behavior.

We explored the effects of the past experiences on both the final target choices and the initial fixations on the contextual cues. To systematically analyze the history effect, we created the general linear model (GLM) that included the indicator variable of mere presence of the specific contextual cues (and its associated targets) and the variable whether the target choice was actually made on this sequence and rewarded (see **Methods**). The variables were created on each past trial between N-1 trial and N-8 trial. Our main interest is whether the rewarded simulations of the event sequence in the past trial, which were the latter variables, biased the sequence selection and simulation in the current trial. The former variables were the control variables that account for the effects that simple exposure to the cue and associated targets on the past trials can have on the current sequence selection and simulation. We hypothesize that the actual simulation of the sequence, which is expressed as the target selection by touch, and experience of complete sequence with contingent reward delivery are necessary to influence the processing of the sequences on the current trial. A mere exposure to the cue and the target would not be enough because it is incomplete and may not be processed as an effective sequence. All past trials of four conditions were included for prediction of a choice of a target, which was associated with a specific contextual cue in a sequence, on a current trial of Competitive cue condition. Note that this analysis is not about a choice of a target *per se* but about the underlying simulation of association sequence that involves both the targets and the associated contextual cues.

First, we examined the patterns of the final target choices that were dependent on the past history of the event sequences. As predicted by our hypothesis, the target choices in the current trials were influenced by the chosen and rewarded cue target sequence. The beta values of indicator variables on each trial are generally positive and decay with increasing trial number into the past (t-tests: N-1 trial, t_16_ = 3.90, p = 0.0013; N-2 trial, t_16_ = 1.47, p = 0.16; N-3 trial, t_16_ = 7.68, p = 9.4^−7^; N-4 trial, t_16_ = 4.15, p = 0.00075; N-5 trial, t_16_ = 3.91, p = 0.0013; N-6 trial, t_16_ = 3.06, p = 0.0074; N-7 trial, t_16_ = 0.87, p = 0.40; N-8 trial, t_16_ = 1.04, p = 0.31) (Fig. 6A; the results were consistent across monkeys, Supplementary Figure 5). The exception was the weaker but still positive effect on N-2. On the other hand, the mere presence of the target and the contextual cue did not have the consistent positive effects on the choices on the current trials.

**Figure 6.**
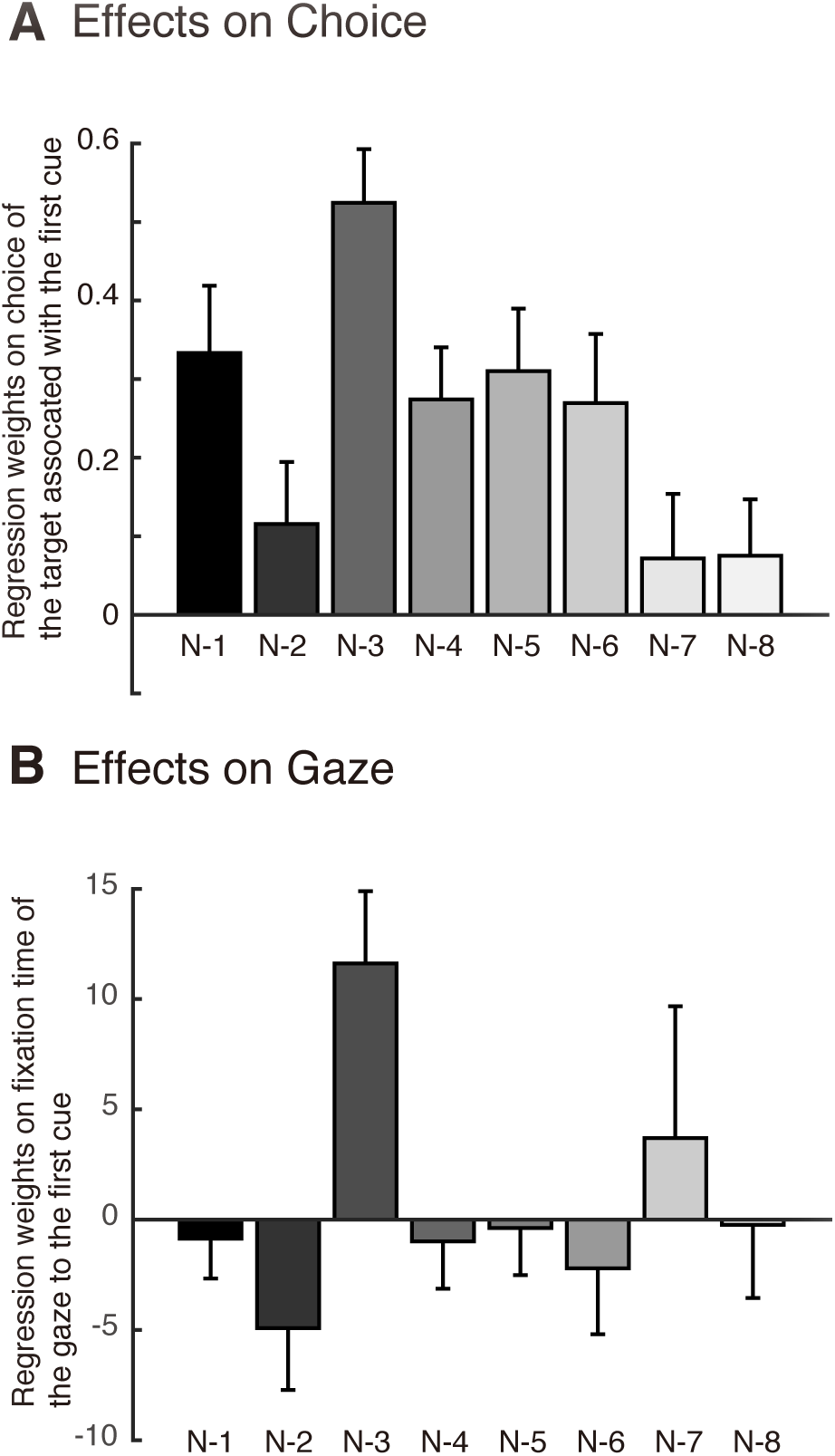
Influences of past episodes of chosen and rewarded sequences on choice and gaze. (A) Beta weights of the individual episodes of the event sequences in past trials (N-1 to N-8) are presented. The event sequence was defined as the sequence of the cue, the touch of the associated target and reward delivery. The choice of target in the current trial was therefore regarded as the choice of this entire sequence and not the target *per se*. The results actually showed the effects of the past episode of the sequence on the choice of the target associated with the same contextual cue that appeared together with the same target, which was then chosen with reward delivery in the past trials. As control regressors, the mere presence of the cue and target sequence was included in GLM. (B) Beta weights of the same GLM on the gaze after onset of the first cue was presented. In contrast the effect on choice, a heterogeneous pattern of past individual episode of sequences was observed with distinct attractive effect of N-3 trial and slightly repulsive effect of N-2.

We also analyzed the effects of the individual past episodes of the cue-target sequences on the patterns of the gaze to the first contextual cue. Here, the predicted variable was the length of the fixation time within a time period. We chose the critical time window from shortly after the onset of the first contextual cue until the 50 ms after the target/second cue onset because it is the time period when relatively pure information processing of the single cue-target sequence of the first cue was possible. The results showed that the effect of the individual past episode of the sequence had heterogeneous effects on the current gaze in this time period. The N-1 trial did not have any strong effect (t-test, t_16_ = -0.48, p = 0.64). However, the N-2 effect had a weak repulsive effect on attention to the same contextual cue in the current trial (t-test, t_16_ = -1.76, p = 0.098) whereas the N-3 episode had an attractive effect on the cue on the current trial (one sample t-test, t_16_ = 3.55, p = 0.0027; the difference between N-2 and N-3: paired t-test, t_16_ = -3.73, p = 0.0018) (Fig. 6B; the results were consistent across monkeys, Supplementary Figure 5). Earlier trials in the past did not have any strong effect (N-4 trial, t_16_ = -0.46, p = 0.65; N-5 trial, t_16_ = -0.18, p = 0.86; N-6 trial, t_16_ = -0.74, p = 0.47; N-7 trial, t_16_ = 0.62, p = 0.54; N-8 trial, t_16_ = -0.071, p = 0.94). The trace of such opposing effects of N-2 and N-3 trials on the current gaze to first cues might mediate the influence of the past episodes of the cue-target sequences on the final choice so that relatively smaller positive effects were observed for the variable of N-2 compared to the effects of the N-3 trial (paired t-test, t_16_ = -4.81, p = 1.9^−4^) (Fig. 6A).

## Discussion

An intrinsic relationship between attention and decision making has been suspected but treated largely as separate entities. Attention in the narrow definition is to seek the relevant part of the sensory information from the environment ^40^. Decision making is to choose the most valuable course of action from alternatives to maximize the reward and minimize the cost. Recent developments have made the definitions less rigid and recognize some interactions between them^1,2,28,29,31,34^. In a natural behavior, these two processes may go hand in hand: the reward objects attract attention and then attention guide the following sequence of the actions to achieve a goal of obtaining the reward^8^. However, it is still not known how such goal-directed operation and attention are operating in a typical decision situation with multiple goals. To examine these mechanisms, we have specifically designed the choice task with sequential events and analyzed gaze patterns. We found that the sequential simulation underlay attention and guided decision process.

Information seeking to reduce uncertainty and reward-guided attention have been treated as separate forces to drive attention^1,5^. However, recent reports suggest that these seemingly discrete influences can be seen as intrinsically connected. Hunt and his colleagues showed bias of information seeking in the situations with more expected reward, which the authors called “approach-induced bias”^9^. Monkeys performing the decision task with advanced information for reward also showed more preference for the information of the greater reward^10^. These results can be interpreted that attention/information seeking is an intrinsic part of a goal-directed operation in foraging behavior so that the properties of the goal itself influences the information seeking behavior^38^. Thus, the current results of sequential gaze patterns are also consistent with this emergent framework of attention and goal-directed behavior.

Attention or gaze patterns observed in previous decision making studies were regarded as random process^28,29^. But the current results suggest alternative interpretations. Specifically, the framework of the sequential simulation suggests that information sampling by gaze and the final choice are connected in the underlying sequential representation. Similar ideas have been proposed as an explanation of biased information seeking behavior in card sorting task performed by human subjects^9^. Our idea is similar in spirit and expresses it as a manifestation of the underlying sequential simulation. A similar idea of simulation and attention have been also proposed in a study of multi-step decision making tasks with examination of gaze patterns^31^.

It might be argued that the monkey subjects in the current study were over-trained and all the gaze patterns and choice behaviors were already habitual and mostly conditioned responses. If that were the case, these monkeys would not be affected by the variability of the local statistics of the event episodes and the recent experiences with those events. In contrast, if the monkeys were still using simulations to guide their behavior, the source of the information for the sequential simulation may be coming from the recent past experiences. Indeed, we found that the past episodes of the sequences exerted significant effects on both the current choices and the patterns of the gazes. These results were reasonable because our data came from the early stages of the training, which was within 1 month of the acquiring the procedure of all conditions. It is also notable that our task involved two actions (two touch targets) in all conditions and two outcomes depending on the choice. Habituation and insensitivity to the recent reward contingency does not usually occur in such situations^41^.

Previous ideas of simulation assumed the Markov property of the individual events, which are independent state spaces^32^. Our idea does not deny some independence of event spaces because the ongoing simulation of event sequence can be flexibly switched to another in the middle of decision making. If it were rigidly chunked as a string of events, the execution the sequence would be automatically completed to the end without any opportunity of the change. Nevertheless, it is not completely flexible and the switching from one sequence to another involved behavioral costs, which was observed in our decision task (Fig. 5), and biases in the studies of information seeking^9^. Related results of the mixture of the heterogeneous types of simulations have also been reported in relation to the successor representation in reinforcement learning framework^26,27^.

Although the current study examined the phenomena of sequential simulation in behavior, the underlying neural mechanisms of sequential simulation can be heterogeneous. Indeed, there have been reports of multiple mechanisms that can support the computations of sequential simulations. In hippocampal formation, Redish and his colleagues reported the phenomenon called vicarious trial and error (VTE) in hippocampal neurons, in which neurons showed the simulation-like activities at the decision point of a maze. Such VTE-like phenomena was also observed in humans performing exploratory behavior and was impaired in amnesic patient with hippocampal lesions^42^. Successor representation proposed in reinforcement learning framework to account for the sequential nature of the event spaces seems to explain hippocampal activity and many of the goal-directed behavior with long sequence of events ^25–27,43^. However, the neural activity patterns related to spatial attention have been reported in dorsal attentional network including the dorsolateral frontal cortex and parietal areas^1,44–46^. There have to be some mediating structures because the hippocampal formation and these brain areas were not connected. It was reported that medial parietal areas receiving inputs from hippocampal formation showed the activity related to cue induced anticipatory spatial attention^47^. Though the current study does not record neural activity and cannot determine the neural mechanisms, it is possible this pathway might contribute to the eye-movements and related simulation/search behavior observed in the present and previous studies^31,32,42,48,49^.

It is known that memory from past episodes can influence attention and decision making. It has been proposed that decision is made by sampling information from the of the past experiences^50^. Recent studies of the foraging decisions also showed the heterogeneous effects of the past experiences^51^. Furthermore, it was also shown that even single events were remembered and had influence on the decision^52^. Such ideas of sampling and replay of the episodic memory on learning and decision making was recently proposed and supported by some experimental evidence^25,52–54^. Thus, the current results of the influences of the past episodes on attention and decision process are consistent with the accumulating evidence and related ideas. But this emerging framework of simulation/replay is still in a very nascent stage. The observations have to be replicated and ideas should be tested rigorously in future studies.

## Methods

### Subjects

Two male monkeys (*Macaca fuscata*) weighing 10.0 kg (*monkey B*) and 9.8 kg (*monkey M)* were used in the current study. The monkeys were cared according to the National Institutes of Health guidelines and the guidelines of the Tokyo Metropolitan Institute of Medical Science. All animal care and experimental procedures were approved by the Animal Care and Use Committee of Tokyo Metropolitan Institute of Medical Science.

### Apparatus

Each monkey sat in a primate chair with its head restrained and with small push buttons (37 mm in diameter) beneath the right hand. The left hand was comfortably restrained. A 19 inch LCD touch screen monitor (TouchTEK, Yokohama, Japan) was placed 21 cm in front of the monkey. We used the TEMPO-NET system (Reflective Computing, Olympia, WA) to control the sequence of visual displays in the behavioral task and the delivery of the reward with opening and closing of a solenoid valve.

### Surgery

For the surgery of head fixation, anesthesia was induced in the monkeys with ketamine hydrochloride (10 mg/kg im) and atropine sulfate. The monkeys then underwent aseptic surgery while anesthetized with pentobarbital sodium (20–25 mg/kg iv), and antibiotics and analgesics were used to prevent infection and pain. During the surgery, polycarbonate and titanium screws were implanted in the skulls of the monkeys, and two plastic pipes were rigidly attached with acrylic resin.

### Task

The monkeys were trained to press a small hold button using the right hand to initiate the trial. After the onset of the fixation point in the center of the screen (white square, 4.9° × 4.9° of visual angle), then the monkey briefly fixated on it. If the monkey continued to fixate on this point for 500 ms, the first contextual cue was presented (a picture of a flower, 9.8° × 9.8° of visual angle) either at the left, right, up or down positions relative the center spot (eccentricity 11.8°). Free eye movements were allowed after this time point while the hold button was kept pressed by the right hand. After 300 ms, the second contextual cues were presented together with action targets (red or green square, 4.9° × 4.9° of visual angle; touch region was set to the size of 9.8° × 9.8° of visual angle to accommodate the inaccuracies of the touch positions), which were presented 90°-rotated positions relative to the contextual cues (if the contextual cues are presented to the left and right positions, the targets are presented to the up and down position and vice versa). In every block of 10-20 trials, the horizontal and vertical positions of the contextual cues and targets are interchanged. Within the block of trials, the positions were pseudo randomly assigned to the contextual cues and the targets. The contextual cue(s) instructed the monkey to reach for the one of the targets. After the presentation of the targets, the monkey was free to release the hold button and reach for the target based on the contextual cues. The release of the hold button extinguished the contextual cues to discourage the further information sampling after the initiation of the reaching movement. In the conditions of Single, Distractor, and Cooperative, the contextual cues determined the rewarding target for 100 % of the times. The touch to this rewarding target led to the delivery of the juice reward (0.6 ml and 0.8 ml for the monkey B and monkey M, respectively) immediately after the touch. In Competitive condition, the reward is delivered at 50% of the times regardless of the specific target touched. The percentages of trials were 40% of all trials for Single condition and 20% of all trials for each of other conditions. The reward was followed by an inter-trial interval of 1 s. The monkeys performed the task highly accurately (about 90% of trials led to a rewarding choice) in the conditions of Single, Distractor, and Cooperative. The monkey could not anticipate the condition of the given trial until the presentation of the second cue. Thus, the optimal strategy for the monkeys was to faithfully process each of the contextual cues.

### Eye movement recording and analysis

The gaze positions were monitored at 1000 Hz with an infrared eye-tracking system (Applied Science Laboratory, Bedford, MA). The online calibration was made according to the positions of the gaze at the time of the target touch based on the fact that the eyes were almost always fixated on the chosen target position. In the off-line analysis, the regions around the targets and contextual cues were carefully chosen based on this eye-hand coordination (a square region of 10.0° x 10.0°). The probabilities of the gaze to each of the targets and the contextual cues were calculated for each sampling time point (1000 Hz) by dividing the number of occasions of the gaze falling on the falling on the defined regions with the number of the total possible instances that these events can occur ^34^. The same calculations were performed for each condition, each case of the cue-choice consistencies (1st-cue consistent choice and 2nd-cue consistent choice), types of the target (chosen/unchosen targets), and types of the contextual cues (choice-consistent/choice-inconsistent cues). Because the patterns of behavior were similar across the monkeys, the data of the sessions across monkeys were entered into the statistical analyses.

### Regression analysis of cue-target sequence selection

To examine the strategy of the monkey in selecting cue-target sequence, we conducted regression analysis of the effects of the trial history. We had explanatory variables of both the mere presence of the cue target pair and interaction terms of previous choice and reward as. They were coded as 1 if a specific cue was the sequence of a contextual cue, a touch of an associated target, and a resultant reward. In other words, the first variables were mere presences of a cue and an associated target regardless whether the pair was actually used to choose the target and the reward is delivered as a result of the choice. The presence or absence of a specific cue-target was coded as 1 or 0. In contrast, the latter variables reflected whether the subject actually used the association to choose the target and received the reward resulting in the complete sequence of cue-target-touch-reward. If a specific cue-target is chosen and not rewarded, they were coded as -1. If the specific cue was not selected in a given previous trial, they were coded as 0 regardless of the reward outcome. The effects of past trials were individually estimated through N-8 trials. With the additional constant term, the beta weights of these trial history effects were estimated.

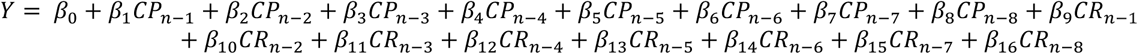

CP is cue-target presence regardless whether it was used to choose the target and CR stands for the cue that is associated with the target of the choice, which in turn led to a reward delivery. We repeated the same procedure for each of the specific contextual cue-target pair and the beta weights were estimated in predicting the subjects’ actual cue-target sequence selections, which was determined by the choice of the target associated with a specific contextual cue. The beta weights 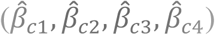 were combined with weighting of the cue-target specific beta values according to their variances 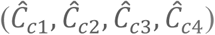 across the specific cue-target associations.

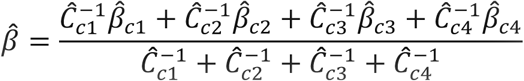

Similar technique of the variance-based weighting was used in a previous study of decision making and credit assignment processes^55^. These procedures were employed for each individual session and obtained values of beta were averaged across sessions.

## Acknowledgements

We thank Yoshihisa Nakayama, Osamu Yokoyama, Tomoko Yamagata, Hiroaki Ishida and Nobuya Sano for technical assistances. R.A. is supported by Research Fellowships for Young Scientists from the Japan Society for the Promotion of Science (JSPS). We thank Nils Kolling, Masaki Isoda, Jill O’Reilly and Joshua Brown for helpful comments and discussions on the previous versions of the manuscripts.

## Author Contributions

R.A. conceived the project, conducted experiments, and analyzed the data. E.H. supervised the project. R.A. and E.H. wrote the paper.

## Additional Information

The authors declare no competing financial interests.

